# The predictive value of double-stranded RNA for A-to-I editing detection

**DOI:** 10.1101/2022.01.29.478304

**Authors:** Guy Shur, Yuval Tamir, Alal Eran

**Author notes:** Equal contributions (Guy Shur, Yuval Tamir).

## Abstract

**Motivation:** Adenosine-to-inosine (A-to-I) RNA editing, a crucial reaction for many processes that contribute to transcriptome plasticity, is both widely common across the transcriptome and difficult to predict due to a lack of distinctive genomic characteristics that can be obtained and analyzed computationally. An exception to this is the secondary structure of RNA molecules, which has been shown to have a major impact on the selectivity and specificity of the enzymes responsible for A-to-I editing. Yet, this information is rarely used for the task of editing site prediction.

**Results:** Here, we demonstrated the value of using base-pairing probabilities of RNA nucleotides to classify genomic sites as A-to-I RNA editing sites, using large-scale truth data which we compiled and make available for use in training future models. Our analysis suggests that the span of four bases from –2 (upstream) to +1 (downstream) of a putative editing site is most informative in this regard. A classifier trained on base-pairing probabilities alone performed with a positive predictive value (PPV) of 0.68, a negative predictive value (NPV) of 0.64, and an area under the receiver operating characteristic curve (AUC) of 0.71. By identifying structure-related features that are informative for detecting A-to-I RNA editing sites and quantifying their predictive value, this work advances our understanding of A-to-I editing determinants.

**Availability:** All source codes and data are available at https://github.com/Ally-s-Lab/P-BEP

## 1. Introduction

RNA editing is the process by which RNA molecules of living organisms undergo intentional modifications during and/or after transcription. The most common form of non-ubiquitous RNA editing in vertebrates is a site-specific process in which an adenosine nucleoside is transformed into inosine via a deamination reaction (Zinshteyn and Nishikura, 2009). This reaction, termed A-to-I editing, plays a key role during development and differentiation, where its spatiotemporally-dynamic nature greatly enhances transcriptome plasticity (Tan *et al*., 2017; Hwang *et al*., 2016; Venø *et al*., 2012; Osenberg *et al*., 2010). In many cases, a typical editing pattern (and sometimes a subsequent shift of that pattern as a tissue matures) is crucial for proper tissue development and function. Such is the case for glutamate ionotropic receptor AMPA type subunit 2 (*GRIA2*) transcripts, in which editing results in a glutamine to arginine recoding in essentially 100% of the transcripts, making the translated AMPA receptor subunit impermeable to Ca^2+^ (Tang *et al*., 2012). Editing at a different site, which recodes an arginine to a glycine, is important for controlling the receptor’s desensitization and recovery rate, and occurs more frequently as the cell matures (Lomeli *et al*., 1994; Wright and Vissel, 2012).

Atypical RNA editing patterns have been associated with numerous diseases and disorders, including cancer, amyotrophic lateral sclerosis, schizophrenia, and autism spectrum disorder (Maas *et al*., 2006; Eisenberg and Levanon, 2018; Paz *et al*., 2007; Eran *et al*., 2013). In light of the severe known and hypothesized implications of altered A-to-I editing, significant efforts have been invested in identifying and characterizing A-to-I editing sites. Accordingly, numerous biochemical and computational methods have been developed for these purposes, and several databases containing millions of editing sites collected from different studies have been compiled (Picardi *et al*., 2017; Ramaswami and Li, 2014; Kiran and Baranov, 2010). Due to the time and cost needed to biochemically identify or verify editing at any given position, the majority of reported sites are based on DNA/RNA mismatches and various predictive models. Given a set of true cases (editing sites) and false cases (non-editing sites), a typical machine-learning model is built by combining often-hidden characteristics of the items in each set that can serve to distinguish items in one set from items found in the other. Although the reported accuracy of such models has dramatically increased in recent years, most train on small datasets (containing up to a few thousand sites) and seemingly study-specific features (Chen, Feng, Yang *et al*., 2016; Zhang and Xiao, 2015; Ouyang *et al*., 2018; Ahmad and Shatabda, 2019; Sun *et al*., 2016). As a result, many of these models perform significantly worse when tested on datasets other than that upon which they were trained, suggesting that they were overfit to the small training data. Hence, to globally improve the accuracy of existing models, it is important to integrate known quantifiable characteristics of A-to I editing sites, and to measure their predictive power on a large dataset.

One of the only known elements common to all A-to-I editing sites is the emergence of a double-stranded structure in the RNA molecule around the editing site, shown to be required for processing by the adenosine deaminase acting on RNA (ADAR) family of proteins, the catalysts of A-to-I RNA editing (Barraud and Allain, 2012; Thomas and Beal, 2017). Indeed, Liu *et al*. exploited the predicted secondary structure in their analysis and reported that RNA structure is one of the key features for predicting A-to-I editing sites (Liu *et al*., 2019). Nonetheless, the double-stranded nature of edited transcripts is not widely used for editing prediction, with examples of incorporating such knowledge into predictive models being rare to non-existent. In comparison, polypeptide secondary structure is commonly used when predicting protein structure and in protein-protein interaction site prediction (Zeng *et al*., 2020; Lee *et al*., 2007; Ofran and Rost, 2007).

Here, we assessed the predictive value of the propensity to form a double-stranded structure around *bona fide* A-to-I editing sites for the purpose of A-to-I RNA editing site detection. By using only the predicted pairing probabilities of bases in the vicinity of a given site, our single feature-based model was able to predict A-to-I editing sites with a PPV of 0.68, NPV of 0.64, and AUC of 0.71. Our work further identified specific positions where base pairing probability is most informative for editing prediction, namely, positions immediately upstream to the editing site, as well as editing sites *per se*. Incorporating these features into future A-to-I RNA editing detection efforts will improve the investigation, and therefore understanding, of this key regulatory mechanism in health and disease.

## 2. Methods

### 2.1 Training and test set compilation

We first compiled a large set of human genomic locations identified as editing sites based on previous studies (**Table S1**). These include 127 biochemically validated sites which we mined from the literature (**Table S2**). We expanded this highest-confidence set to include 462,067 sites identified by large-scale editing detection efforts (Porath *et al*., 2014; Sakurai *et al*., 2010; Levanon *et al*., 2005), all included in RADAR, and 317,552 sites supported by at least 100 reads from REDIPortal, an online database of editing sites (Picardi *et al*., 2017). We converted sites reported in hg19 coordinates, including the entirety of the REDIPortal dataset, to their respective GRCh38 coordinates using LiftOver (Kent *et al*., 2002) or Crossmap (Zhao *et al*., 2014). To avoid false positives, we filtered out sites that overlap genomic A>G and T>C SNPs detected in 71,702 whole genomes of gnomAD V3.0 (Karczewski *et al*., 2020). We also removed sites within 5bp of gnomAD V3.0 indels, because indel-containing reads are often misaligned. In total, 353,020 sites from three high-confidence profiling studies and 242,700 sites from REDIPortal remained, for a total of 355,276 unique true positive (TP) training sites. For true negatives (TN), we used common genomic A>G SNPs. Specifically, our TN training set included 293,848 genomic SNPs with minor allele frequency (MAF) > 0.1 in 71,702 gnomAD V3.0 genomes.

For testing, we used an independent set of 3,235 high-confidence TP editing sites (Tran SS, Jun HI, Bahn JH, et al.,2019), and 3,234 randomly selected TN gnomAD genomic A>G SNPs with MAF > 0.1, that have no overlap with the training sites.

### 2.2 Quantifying dsRNA formation probabilities

To assess the predictive power of a double stranded structure for A-to-I editing detection, we focused on nucleotide-level base pairing propensity. Specifically, we examined the pairing probabilities of each nucleotide within 25nt of a site (i.e. 25nt up- and 25nt downstream, a total of 51nt). For that purpose, we used the ViennaRNA RNAplfold program with the -p option to predict sequence folding, treating each sequence as if it were transcribed directly (Lorenz *et al*., 2011). For each nucleotide within 25nt of the editing site, we used a sliding window of 70nt to calculate the probability that it is unpaired, and reported its negation.

Toward that goal, we identified and imported the surrounding sequence of each site in the training and testing sets using the Biopython package (Cock *et al*., 2009). Namely, for each site S, we fetched the 191bp genomic interval ranging from 70+25 upstream to 25+70 downstream of S. That interval served as input to ViennaFold RNAplfold, executed as described above, which in turn yielded a vector of probabilities of length 51, corresponding to the base pairing probability of each nucleotide within 25nt of S (Figure 1A). These probabilities serve as features for modeling the predictive value of dsRNA for A-to-I editing in cis.

**Fig 1.**
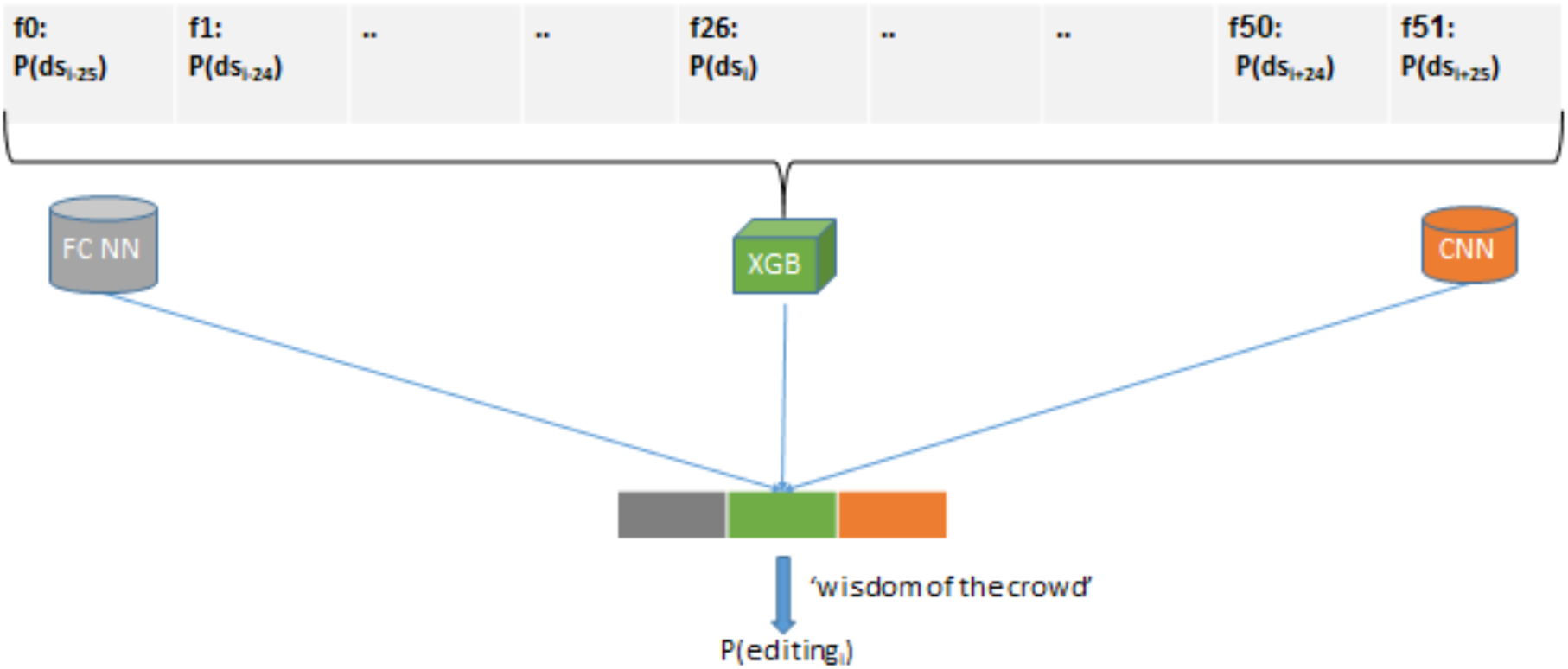
Schematic depiction of the model used to predict A-to-I editing sites. The input vector comprises 51 features (see section 2.2). Edit site probability is calculated by three different classifiers (XGB, fully-connected NN, convolutional NN), and the average value of all probabilities serves as the final prediction.

### 2.3 Model architecture

We trained three independent classification frameworks to predict A-to-I RNA editing at a site, based on base pairing probabilities of its vicinity, namely nucleotides ≤25nt away. Namely, for each site For testing, each model made a prediction on an out-of-sample test set. A general prediction was then made based on the mean prediction of the classifiers (“wisdom of the crowd”). The general pipeline employed is described in fig 2.2. The reason the architecture depicted was chosen is that empirical results suggest this was arrangement was more robust and less overfitting when examined with test data. Each model was trained with 10-fold cross-validation on the entire training set, and subsequently tested on an independent test set.

**Fig 2.**
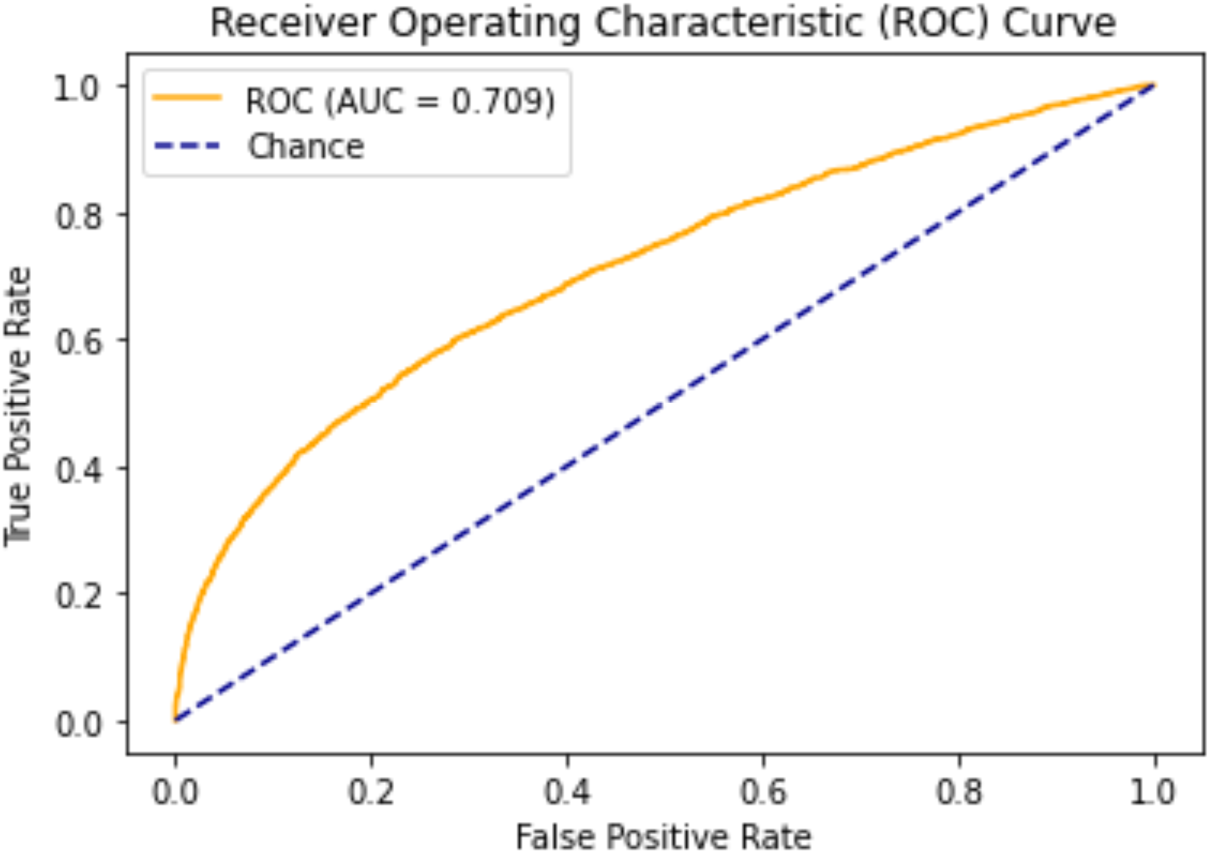
Model performance. Shown is the receiver operating characteristic (ROC) curve. The area under this curve (AUC) is 0.71.

**Fig 3:**
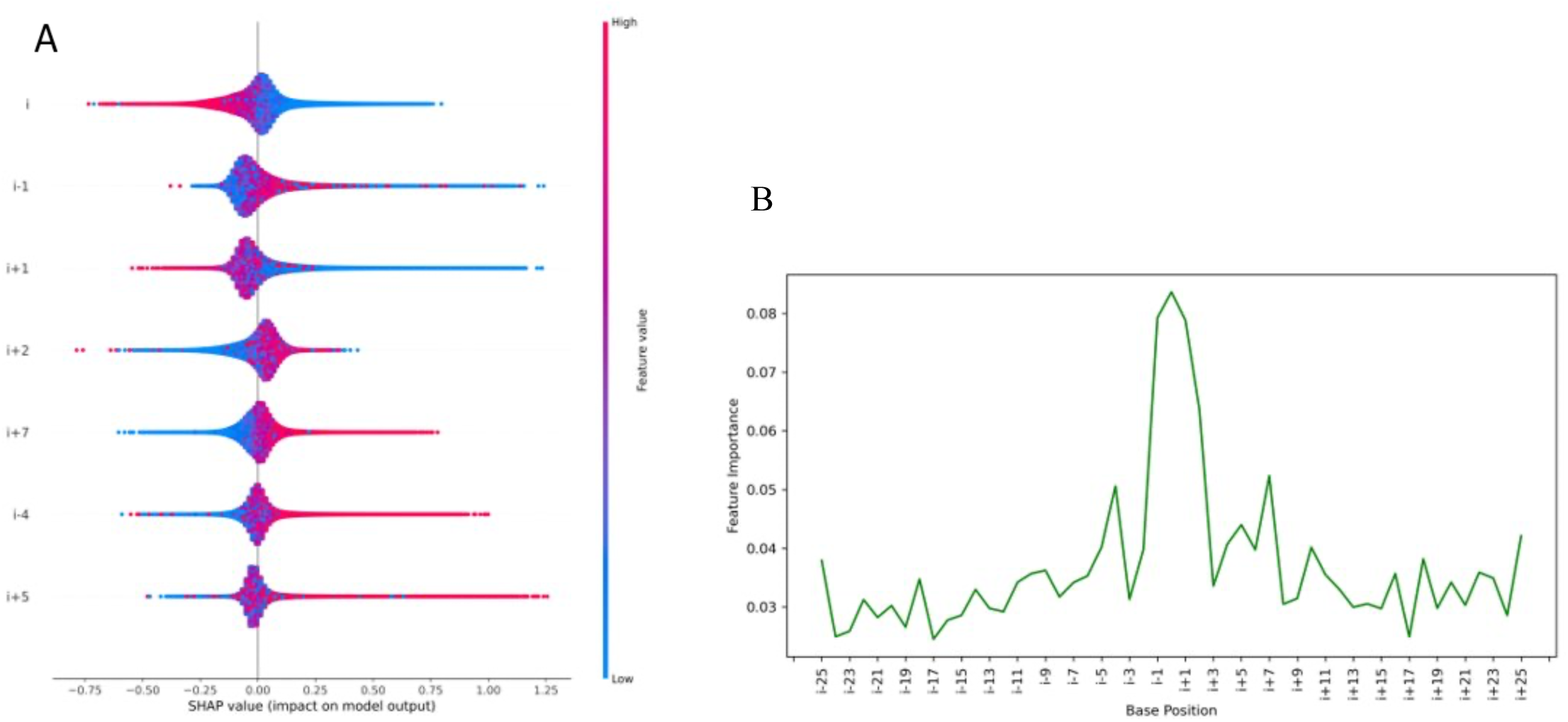
**(A)** Shapley-values summary plot. The summary plot combines feature importance and feature effect; *Feature importance:* Variables are ranked in descending order, with feature indices counting upstream to downstream. *Feature effect:* The horizontal lines correspond to the impact on the decision. (**B**) Distribution of the mean-absolute Shapley-value of bases in the sequence.

### 2.4 Evaluation metrics

To evaluate the performance of our model, four evaluation metrics are used in this study: accuracy (ACC), precision, recall, F-measure, and area under the receiver operator characteristic curve (AUC).

### 2.5 Classifier implementation

Our deep learning framework was implemented with PyTorch (http://pytorch.org/), a popular deep learning package developed by Facebook. XGBoost was implemented with the XGBoost python package (https://xgboost.readthedocs.io/en/latest/python/index.html). The loss functions that were used were Logistic:binary for XGB and CrossEntropyLoss for FC-NN and CNN. Logistic:binary outputs a single value, which predicts the most likely class upon activation. CrossEntropyLoss outputs a vector of length K, corresponding to K classes. The highest value index of the output is the predicted class. The optimizer used for all NNs was Adaptive Momentum (Adam), with a learning rate of 1e-4.

## 3. Results

### 3.1 A collection of large-scale training data

We assembled and formatted a collection of 355,276 previously published editing sites and 293,848 known genomic SNPs as training samples representing true positives and true negatives, respectively. Our true positive set includes subsets of higher confidence (127 experimentally validated or evolutionarily conserved sites, and 3,564 sites discovered using a biochemical approach, ICE-seq. We provide our entire training set for use by the scientific community in future analyses as a possible benchmark set, which may prevent overfitting issues that arise from the use of small training sets.

### 3.2 Model evaluation

All scores are reported on an independent out-of-sample test set comprising 6,469 sites, as described in section 2.1.

**Table 1:** The scores of chosen metrics, as described in section 2.4, on the test data with the label 1 corresponding to an editing site and the label 0 corresponding to a non-edited base.

|  | Precision | Recall | F1-Measures |
| --- | --- | --- | --- |
| Label 0 : | 0.64 | 0.72 | 0.69 |
| Label 1 : | 0.68 | 0.6 | 0.65 |
| Accuracy |  |  | 0.66 |

### 3.3 Model inference

We used the Shapley value-based python package ‘shap’ (https://github.com/slundberg/shap) to visualize the impact of the most informative bases with regard to a prediction (Lundberg and Lee, 2017; Lundberg *et al*., 2020). Essentially, the Shapley value is the average expected marginal contribution of one feature after all possible combinations have been considered.

## 4. Discussion

A-to-I editing prediction is difficult due to the strikingly low information content of the genomic sequence around an editing site. This difficulty has led researchers to develop various models that seek to tackle the problem. Pinto and Levanon (2019) compiled a list of best practices and points to consider when attempting to predict A-to-I editing sites. Shu *et al*. (2014) created a model based on several dozen pre-selected features. Here, our objective was to verify and quantify information attained by analyzing the predicted pairing probabilities of bases in RNA molecules alone for the purpose of A-to-I editing site prediction only by analyzing the predicted base pairing probabilities of RNA molecules. Evaluation of our meta-predictor showed that this data can be used to effectively distinguish between editing sites and controls. We next sought to identify the most informative positions, relative to a putative editing site, using SHAP analysis to not only determine these positions but also to assess how they affect the prediction. The results suggest that those bases -2 downstream to +1 upstream to the site of editing are the most informative and display a propensity to be unpaired, with the nucleotide immediately upstream of the editing site being the most informative. This result supports reports that bases immediately neighboring editing sites are slightly enriched with specific nucleotides (Nigita *et al*., 2015). Notably, both high- and low-pairing probabilities of certain sites are indicative of the presence of an editing site. The enrichment of unpaired bases at the editing site could be explained by enhanced accessibility of the edited adenosine, and in some cases (such as the aforementioned GLuR-2 R/G editing site), editing replaces an A-C mismatch with an I-C pair, thereby increasing secondary structure stability(Barraud and Allain, 2012). Furthermore, 9 of the 10 most informative bases are found at most 6 positions away from the presumed editing site, also in line with our expectations, given that this area has been shown to be in direct interaction with human ADAR2 using X-ray crystallography (Thomas and Beal, 2017).

Due to the low number of studies implementing RNA structure related data, much of our workflow was based on experimentation. The window size of 51 bases (25 bases on either side of the editing, together with the site itself) used in the analysis was selected on the basis of trial and error, and as a result, supports previous findings on the most informative span of bases (Chen, Feng, Ding *et al*., 2016; Xiao *et al*., 2018).

It should be noted that our decision to predict RNA folding on the direct basis of the DNA sequence, which corresponds to a folding prediction of unedited and not mature RNA, may not be applicable in cases where other editing events previously occurred. Moreover, many sites in the true positive set have not been fully validated as *bona fide* A-to-I editing sites. Overall, our results suggest that the information gain from RNA secondary structure is significant and that predicted base pairing probabilities or an equivalent metric can and should be used as a primary feature in future A-to-I editing site predictive models.

## Notes

https://github.com/Ally-s-Lab/P-BEP

### Competing Interest Statement

The authors have declared no competing interest.

